# The supramolecular landscape of growing human axons

**DOI:** 10.1101/2020.07.23.216622

**Authors:** Patrick C. Hoffmann, Stefano L. Giandomenico, Iva Ganeva, Michael R. Wozny, Magdalena Sutcliffe, Wanda Kukulski, Madeline A. Lancaster

## Abstract

During brain development, human axons must extend over great distances in a relatively short amount of time. How the subcellular architecture of the growing axon sustains the requirements for such rapid build-up of cellular constituents has remained elusive. Human axons have been particularly inaccessible to imaging at molecular resolution in a near-native context. Here we apply cryo-correlative light microscopy and electron tomography to growing axonal tracts from human cerebral organoids. Our data reveal a wealth of structural details on the arrangement of macromolecules, cytoskeletal components, and organelles in elongating axon shafts. In particular, the intricate shape of the endoplasmic reticulum is consistent with its role in fulfilling the high demand for lipid biosynthesis to support growth. Furthermore, the scarcity of ribosomes within the growing shaft suggests limited translational competence during expansion of this compartment. These data provide an unprecedented resource and reveal a molecular architecture that helps explain the unique biology of growing human axons.

## Introduction

Mammalian neurons are highly unusual cells. With an elaborate tree-like structure and elongated shape sometimes up to a meter in length in adult humans, they have the highest surface area to volume ratio of any cell in the body (*1*). Neurons are also extremely polarized, with structurally and functionally distinct axon and dendrite compartments (*2*, *3*). While dendrites receive signals, the axon transmits the electrical signal to downstream targets. It is this directional relay, together with the specific topology of axonal connections and their bundling in defined tracts, that allows for information transfer, and ultimately cognition. The formation of such long-range tracts depends upon the coordinated growth of thousands and even millions of immature axons, sometimes over great distances.

How the axon achieves such remarkable elongation remains to be determined, but its subcellular architecture is likely to reflect unique requirements for growth. For example, the organelle composition of the developing axon must be such that the neuron, without going through cell division, can undergo size increase rates akin to rapidly dividing cancerous cells (*4*). How is the cytoskeleton organized within growing axons? How is the constant supply of lipids and proteins, necessary for the large increase in surface area, ensured? How does the neuron deal with the challenge of molecular transport within the rapidly extending axon? These are some of the outstanding questions that molecular-resolution insights could help to address.

The cellular organization of different neuronal compartments, including the axon, has been studied by electron microscopy (EM) since its earliest days on fixed nervous tissue from rodents (*5*, *6*). While these studies revealed many ultrastructural features, the resolution and interpretability of classical EM methods are restricted by limited sample preservation. More recently, cryo-EM has enabled analysis of mammalian neuronal cells preserved to the molecular level, thereby visualizing the spatial organization of macromolecular structures in their native environment (*7*–*9*). Nevertheless, it remains challenging to unequivocally identify axons in primary cultures (*2*, *3*) because the cells are dissociated from their tissue context, and therefore exhibit intermixed axons and dendrites. Ultrastructural analysis at a molecular detail-level of human axons within the physiological context of a tract would be invaluable for understanding neurodevelopment, nerve damage, and regeneration (*10*).

Cerebral organoids provide the opportunity to study aspects of human neuronal physiology within the context of a complex 3D cellular milieu mimicking the architecture of the developing brain (*11*–*14*). Recently, we developed an optimized air-liquid interface slice culture paradigm that enables long-term culture of cerebral organoids and promotes the establishment of axon tracts, consisting of dozens to hundreds or even thousands of individual axons bundled together, able to form functional connections (*15*). While the cell bodies are part of the organoid tissue, these tracts project over long distances of several millimeters from the organoid, effectively segregating the axons from dendrites and somas. This feature, combined with the ability to derive cerebral organoids from human cells, makes this method a promising route to examining the cell biology and molecular structure of human axons.

Here, we combine air-liquid interface cerebral organoid (ALI-CO) culture with correlative light and electron cryo-microscopy (cryo-CLEM) to study the molecular architecture of developing human axons in a context closely mimicking *in vivo*. Cryo-CLEM offers unambiguous identification of fluorescent cells, or individual cellular compartments, combined with high-resolution EM imaging of the same, near-natively preserved sample to reveal its underlying molecular structure (*16*). We apply this technology to observe the behavior of growing human axons within tracts in real time followed by electron cryo-tomography (cryo-ET) of the same axons.

## Results

We first examined in more detail axon growth behaviors of ALI-COs. Thick tracts that exited the ALI-COs stained positive for the pan-axonal marker SMI312 and negative for the dendritic marker MAP2 (Fig. 1A and Fig. S1). This, together with their length and physical distance from neuronal cell bodies within the organoid, suggested that these tracts comprise exclusively axons (Fig. 1A and Fig. S1). To monitor growth, we expressed a farnesylated membrane-targeted GFP (fGFP) (*12*, *15*) (Fig. S2) in a subset of cells within the organoid slices. As a result, tracts contained a mixture of unlabeled and fluorescently labeled axons, which extended rapidly within the tracts (Fig. 1B and Movie S1). We measured the extension speed for individual fluorescently labeled axons over several hours to be an average of 691 nm/min, with a peak pace of nearly 3000 nm/min (Fig 1C). Intrigued by the rapid extension rates, we set out to examine the subcellular organization that underlies this unique cellular behavior.

**Fig. 1.**
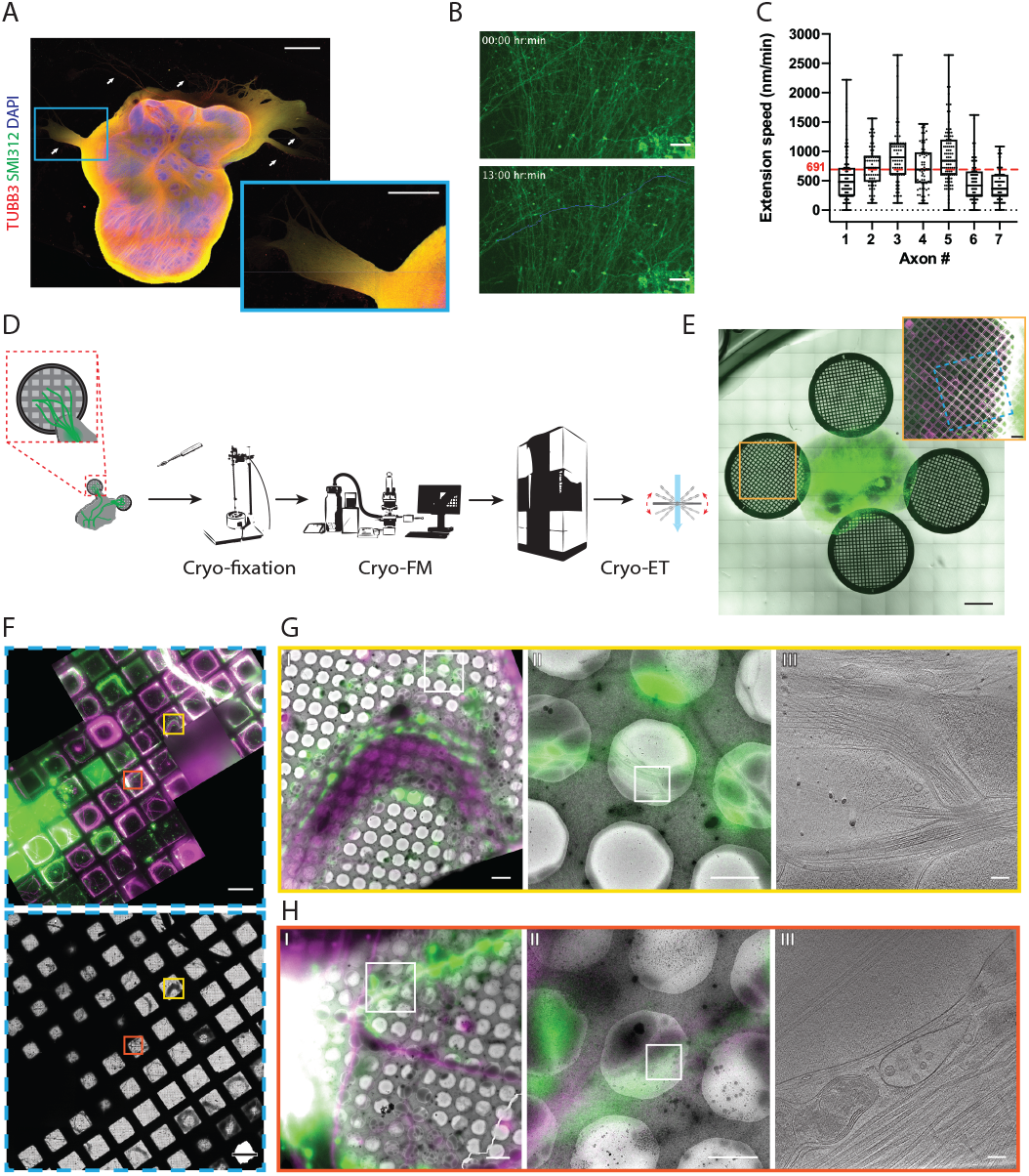
Targeting fluorescently labeled axons within tracts from human cerebral organoids by correlative light and electron cryo-microscopy (cryo-CLEM). (**A**) Immunofluorescence of an air-liquid interface cerebral organoid (ALI-CO) stained for the pan-neuronal marker TUBB3 and the pan-axonal marker SMI312. The box highlights a magnified view of escaping axon tracts. White arrows point to bundles of varying thickness. (**B**) Representative fluorescence microscopy (FM) images of extending fGFP^+^ axons. The top panel is the first frame and the bottom panel is the last frame of the 13 hr-long live FM Movie S1. The images were used to track axon number 4 in (C), and the blue line marks the trajectory of its growth throughout the movie. (**C**) Box and whisker plot of axon extension speed measurements reporting the median, first and third quartiles, minimum and maximum. The individual data points represent the extension speed calculated between two consecutive frames. The measurements were done on 7 axons from 3 different ALI-COs. (**D**) Schematic preparation of ALI-COs for cryo-CLEM including electron cryo-tomography (cryo-ET). (**E**) Overlay of fluorescence and transmitted light overview images of grids placed around an ALI-CO. Lower magnification image shows day 1. Inset shows axons labeled with fGFP and additional tracts stained with SiR-tubulin immediately prior to cryo-fixation after 11-days of growth on the grid to the left of the ALI-CO, indicated by the orange square. (**F**) Cryo-FM (top) and cryo-EM (bottom) overview of the blue area indicated on the grid shown in the inset in (E). Yellow and red boxes indicate the areas shown in (G) and (H), respectively. (**G**) and (**H**) Correlated cryo-FM and cryo-ET on two different grid squares. Subpanels I and II: Overlays of cryo-FM and cryo-EM at different zooms to identify individual fluorescent axons. Subpanels III: Virtual slices through cryo-tomograms acquired at the positions of squares indicated in subpanels II. Scale bars: 1 mm and 500 μm (inset) (A), 50 μm in (B), 1 mm and 200 μm (inset) in (E), 100 μm in (F) 5 μm, 2 μm and 100 nm in (G) and (H) (left to right).

For this, we developed a workflow to prepare axon tracts for cryo-EM (Fig. 1D and S3A). We placed coated EM support grids in close proximity to ALI-COs on organotypic cell culture inserts (Fig 1E and S3B). The signal of fGFP^+^ axons was used to track the behavior of axons over time by live fluorescence microscopy (live-FM). We monitored their growth over the course of 11 days (Fig. 1E and Movie S2). Prior to cryo-fixation, we applied SiR-tubulin to visualize all tracts on the grid (Fig. 1E inset and S3C). Grids on which axon tracts approached the center were detached from the organoid slice using a biopsy punch and immediately cryo-fixed by plunge-freezing (Fig. 1D and S3A). Comparison of the number and position of tracts in FM images immediately prior to cryo-fixation and in corresponding cryo-EM grid overview images indicated that this procedure resulted in good preservation of axon tracts (Fig. S3D). The level of contrast in overview images of individual grid squares suggested that most tracts were sufficiently thin for cryo-ET (Fig. S3D).

To specifically target fGFP^+^ axons within tracts we used cryo-CLEM. We introduced a step of cryo-fluorescence microscopy (cryo-FM) imaging prior to cryo-ET and imaged the center of the same grid by cryo-FM and cryo-EM (Fig. 1F). Fluorescent signals of individual tracts could be correlated to cryo-EM overviews by using landmark features on the grid (Fig. 1G and H, I). This allowed us to distinguish within the tracts fGFP^+^ axons from axons that were positive only for SiR-tubulin (Fig. 1G and H, II). We then imaged fGFP^+^ axons by cryo-ET (Fig. 1G and H, III). By targeting specific segments of individual axons within axon tracts, we were able to establish a direct link between cellular behavior observed live and cellular ultrastructure (Movie S3).

In our cryo-ET data, cellular structures such as protein assemblies and membranes were preserved at molecular detail (Fig. 2A). We observed parallel, unbranched actin filaments, recognizable by their characteristic thickness of 7-9 nm and a subunit repeat of 5-6 nm (*17*) (Fig. 2A, red arrows). Microtubules revealed their individual protofilaments (Fig. 2A, magenta arrows), as well as numerous intraluminal protein densities (*18*). Microtubule bundles were often so dense as to seemingly pose constraints on microtubule-based transport. We observed mitochondria squeezed within these bundles (Fig. 2A, green arrows), suggesting that dynamic rearrangements are required to allow sufficient space for vesicles and organelles to pass. We also identified intermediate filaments about 10 nm in diameter (Fig. 2A, brown arrows). These dimensions match those of neurofilaments (NFs) such as NF-L, NF-M and NF-H (*19*), which serve as signature markers for axons. Our tomograms also revealed coated and uncoated vesicles of variable size (Fig. 2A, yellow arrows, and Fig. S4). Furthermore, we frequently observed large expanses of endoplasmic reticulum (ER) cisternae, which extended into thin membrane tubules tightly associated with microtubules (Fig. 2A, cyan arrows). We also observed a variety of membrane contact sites between ER-mitochondria and ER-plasma membrane (Fig. 2B, white arrows). Finally, as each tract contained several axons, our tomograms frequently visualized axon-axon contacts, a unique physiological feature of axons in their normal tissue context (Fig. 2B, blue arrows). In order to highlight the ultrastructural complexity, we segmented key cellular elements of a fGFP^+^ axon volume (Fig. 2B, bottom panel and Movie S3).

**Fig. 2.**
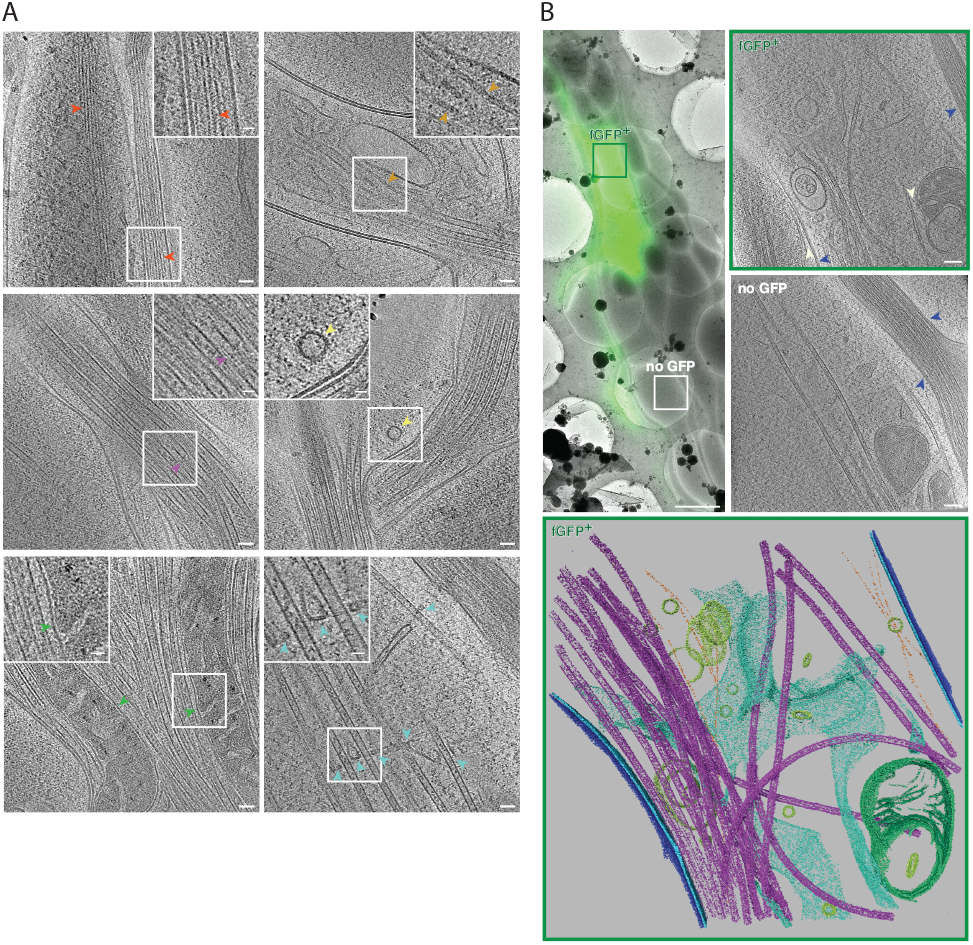
Cryo-ET reveals structural features of developing human axon tracts at molecular resolution. (**A**) Tomograms reveal bundles of unbranched parallel actin filaments, recognizable by actin subunit arrangement (red arrow), microtubules filled with lumenal densities, microtubule protofilaments (purple arrows), mitochondria (green arrows) embedded in microtubule bundles, neurofilaments (brown arrows), vesicles carrying protein cargo (yellow arrow), endoplasmic reticulum (cyan arrows) closely associated with microtubules. Insets show magnified views of the boxed areas. (**B**) fGFP^+^ and untransfected axons are found within the same axon tract, allowing direct phenotype comparison. Top left panel: Overlay of cryo-FM and cryo-EM overview images. Two top right panels: Virtual slices through cryo-tomograms of fGFP+ and control axon. White arrows indicate contacts between ER and mitochondria as well as between ER and plasma membrane, blue arrows indicate cell-cell contacts between different axons. Lower panel: Segmentation model of the fGFP^+^ axon shown in (B), illustrating the complexity of the cellular ultrastructure. Microtubules are shown in magenta, actin filaments in orange, endoplasmic reticulum in cyan, vesicles and other membrane compartments in yellow, mitochondrial membranes in green and the plasma membranes of the fGFP^+^ axon and neighboring axons in blue and dark blue, respectively. Scale bars: 50 nm (20 nm for insets) in (A), 2 μm and 100 nm in (B), top left and top right two panels, respectively.

The 38 tomograms we collected comprised unlabeled axons as well as axons overexpressing either fGFP, GFP-tagged human L1 cell adhesion molecule (L1CAM) or GFP-tagged extended-synaptotagmin isoform 1 (ESYT1) (Fig. S2 and S4). We overexpressed L1CAM, which is involved in axon-axon contacts (*20*), and ESYT1, involved in lipid metabolism (*21*), as possible regulators of tract formation and axon elongation. Because only a subset of cells expressed the GFP constructs, most tracts contained a large proportion of unlabeled, GFP negative axons (Fig. 2B, top left panel). Thus, control axons could be imaged on the same EM grid (Fig. 2B, top right panels), not only simplifying the acquisition of control data but also making comparison more reliable by removing grid-to-grid variability as a source of unspecific differences. We did not observe any obvious structural differences between fGFP^+^ and control axons, nor any notable changes in subcellular structure upon L1CAM-GFP or ESYT1-GFP overexpression (Fig. S4). Taken together, these results indicate that our approach allows one to assess the impact of gene function on molecular cell structure without technical knock-on effects.

A defining feature of axon identity is the parallel arrangement of bundled microtubules (*22*, *23*). We therefore set out to determine the polarity of individual microtubules in tomograms using subtomogram averaging (Fig. 3A). We used axial views of each microtubule average to determine the handedness and thus polarity of the microtubule based on the tilt of its protofilaments (*24*) (Fig. 3A and B). This analysis showed that the majority of bundles had a uniform, parallel microtubule arrangement, further confirming axon identity (Fig. 3C).

**Fig. 3.**
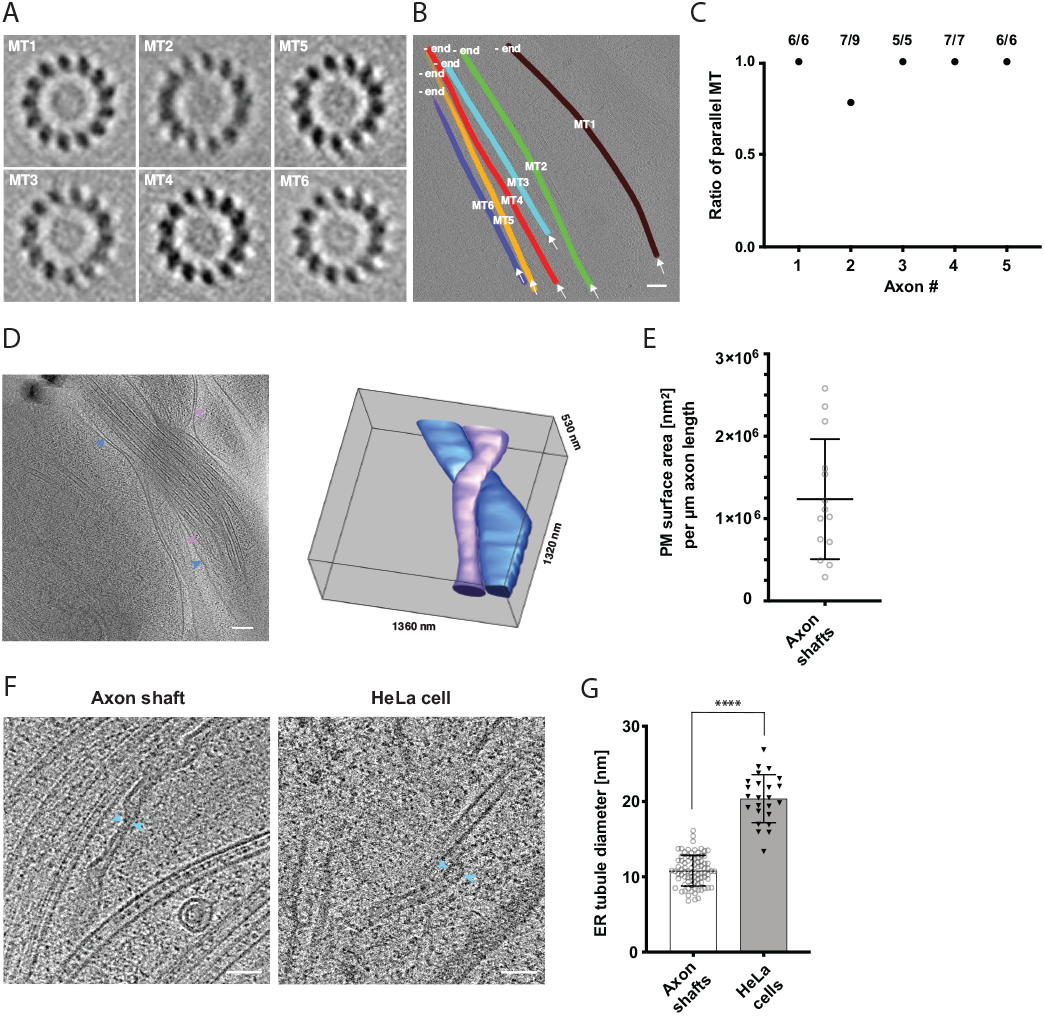
Cytoskeletal and membrane architecture of developing axon shafts. (**A**) Axial views of subtomogram averages reveal the polarity of each microtubule shown in (B). The directionality of the microtubule was determined from the radial tilt of the protofilaments seen in axial views (*24*). (**B**) Tomographic slice with six microtubules depicted as differently colored tubes (MT1-MT6) (same tomogram as shown in Fig 2B, ‘no GFP’ panel). Arrows indicate the viewing direction of the axial views of subtomogram averages shown in (B). The -end of each microtubule is indicated. (**C**) Ratio of parallel microtubules determined by subtomogram averaging in 5 different axons (6-9 individual microtubules per axon). Axon 1 is depicted in (A) and (B). **(D**) Virtual tomographic slice and the corresponding segmentation model of the plasma membrane (PM) of two individual axon shafts. Blue and magenta arrows indicate PM segments visible in the virtual tomographic slice. **(E**) PM surface area measurements in nm^2^ normalized to the length in μm of 14 axon shafts captured in 12 tomograms. (**F**) The curvature of ER tubules is higher in axon shafts than in HeLa cells. Arrows indicate the shortest distance between the membrane bilayer cross sections. (**G**) Diameters of the thinnest ER tubules measured in axon shafts and in HeLa cells (82 and 24 measurements, respectively). Welch’s t test was employed for statistical analysis: p < 0.0001. Scale bars: 100 nm in (A) and (D), 50 nm in (F).

The rapid growth of developing axon tracts must be supported by synthesis of new biomolecules. At average growth rates of 691 nm/min (Fig. 1C), the demand of membrane and lipid molecules must be high. Using the plasma membrane as proxy to estimate the amount of newly synthesized membrane, we segmented the plasma membrane in tomograms and measured the surface area of individual axon shafts (Fig. 3D). Each micrometer of length corresponded to 1.24 × 10^6^ nm^2^ of plasma membrane area on average (SD = 7.3 × 10^5^ nm^2^, N = 14) (Fig. 3E). The broad distribution reflects the variability between the thinnest parts of axon shafts and axonal varicosities. Because each μm^2^ of membrane contains approximately 5×10^6^ lipids (*25*), corresponding to 5 lipid molecules/nm^2^, we estimated the lipid supply to the plasma membrane required to sustain the average growth rate of an axon as approximately 4.3 × 10^6^ lipid molecules/min. This massive influx of new phospholipids into the plasma membrane points to a unique requirement for lipid biosynthesis and transfer.

The ER is the major organelle for lipid biosynthesis. We therefore examined ER ultrastructure to assess whether it may help explain how the unique lipid requirements of the growing axon are supported. Large flattened cisternae (Fig 2B) were reminiscent of the ER observed in cultured neurons (*26*). We further observed tubular segments of axonal ER that were remarkably narrow (Fig. 3F). At their thinnest outer diameter, the ER tubules in axon shafts measured on average 10.8 nm (SD 2.0 nm, 82 measurements). For comparison we measured the thinnest ER tubules in HeLa cells and found them to be about twice the diameter (20.4 nm, SD 3.2 nm, 24 measurements) (Fig. 3F, G). Some of the axonal ER tubules had a local outer diameter of less than 10 nm. Considering that this measurement includes the bilayer thickness, this implies that these ER tubules contain hardly any lumenal space, likely posing constraints on diffusion of ER proteins. The observation that the ER is depleted of lumen whilst adopting highly curved tubular shapes indicates high local membrane surface-to-ER volume ratios, and suggests that the axonal ER structure may be a consequence of maximized synthesis of lipids, produced to sustain high growth rates during axon lengthening.

We anticipated the extending axon to require not only lipids but also proteins for maintaining functionality during elongation. We thus examined the presence of protein synthesis machinery in the elongating axon shafts. While it is known that mature axons do not display extensive Nissl bodies, indicating low ribosomal RNA content (*27*), we reasoned that developing axons, due to their growth state, could have different protein biosynthesis requirements and hence composition. Furthermore, cellular cryo-ET provides the resolution to detect ribosomes, and to even distinguish between monosomes and polysomes (*28*), making it a powerful method for direct detection of the protein synthesis machinery. It was therefore noteworthy that we observed a scarcity of ribosomes within our tomograms of growing axon shafts. In contrast, we readily identified a large number of ribosomes based on their size, shape and high contrast in tomograms of HeLa cells (*29*) and of other neuronal processes (Fig. 4A). In order to assess the occurrence of ribosomes quantitatively, we counted all ribosome-like particles, both cytosolic and membrane-bound, and found on average 2 particles per μm^3^ in axon shafts (SD = 3 particles, n=31 tomograms), 589 particles per μm^3^ in other neuronal processes (SD = 344 particles, n=4 tomograms), and 2314 particles per μm^3^ in HeLa cells (SD = 977 particles, n=5 tomograms) (Fig. 4B). The local ribosome concentration in axon shafts is thus less than 3.5 nM, about 1000-fold lower than in HeLa cells. Of the ribosome-like particles found in tomograms of axon shafts, only a minute number were attached to the ER (Fig. 4A), supporting the idea that the large amounts of axonal ER have a primary function in lipid metabolism rather than protein synthesis. To validate these findings, we tested for the presence of five distinct ribosomal proteins in neurons from dissociated organoids by immunofluorescence (Fig. 4C and S5). In agreement with the cryo-ET data, axon shafts identified as SMI312^+^/MAP2^−^ showed significantly lower ribosomal signal than dendrites (SMI312^−^/MAP2^+^) (Fig. 4D).

**Fig. 4.**
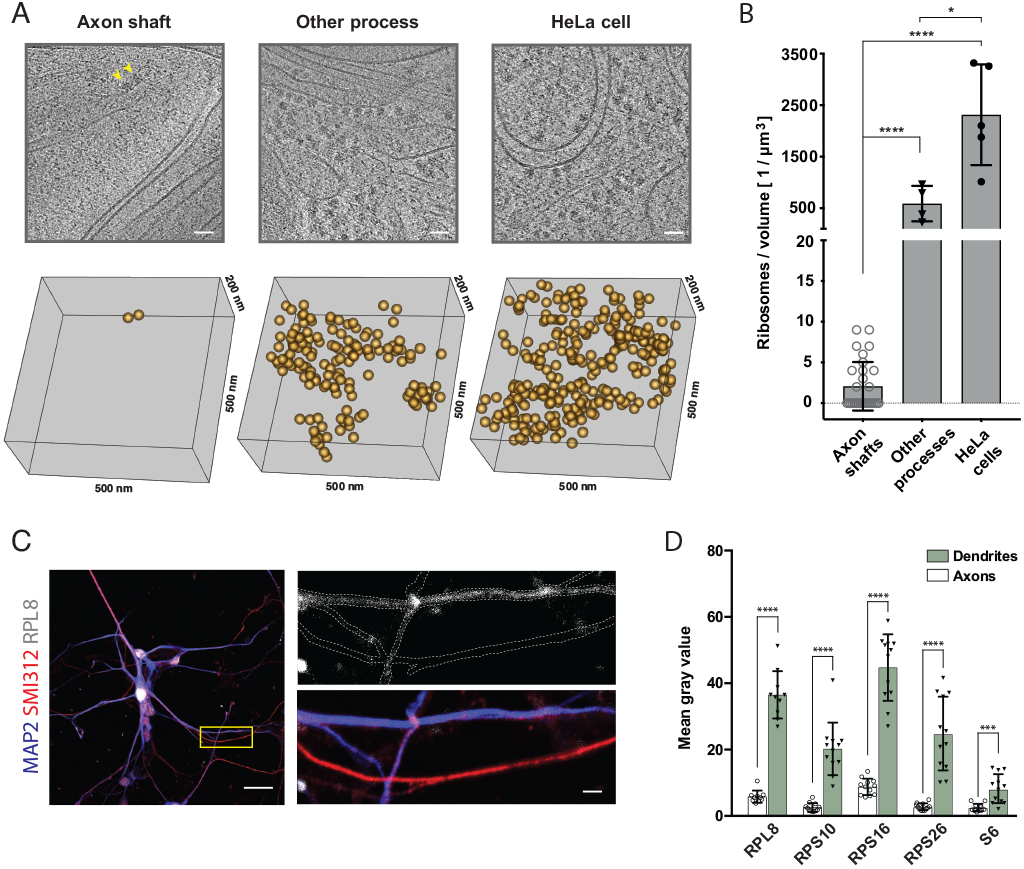
The shaft of the growing axon is scarcely populated by ribosomes. (**A**) Examples of ribosomes observed in cryo-tomograms of axon shafts from ALI-COs, of other cellular processes from ALI-COs, and of HeLa cells. The bottom panel shows 0.05 μm^3^ cryo-ET volumes, corresponding to the area shown in the upper panel. Positions of all ribosome-like particles observed in that volume are shown as orange spheres. **(B**) Comparison of the numbers of ribosome-like particles, normalized to the tomographic volume, observed in axon shafts, other processes and HeLa cells. Individual data points represent individual cryo-tomograms (from 30, 4 and 5 tomograms, respectively). Mann-Whitney tests were employed for statistical analysis: p<0.0001 (****); p<0.05(*). **(C**) Immunofluorescence images of dissociated neurons from organoids reveal low signal for the ribosomal 60S component RPL8 in axon processes (identified by SMI312^+^/MAP2^−^ labeling) in comparison to dendrites (identified by SMI312^−^/MAP2^+^ labeling). The yellow box outlines the area magnified in the right panel. The top image of the right panel shows the immunofluorescence signal for the ribosomal subunit RPL8. The bottom image shows the SMI312/MAP2/RPL8 composite. The white dashed line depicts the outline of axons and dendrites and was traced based on the MAP2 and SMI312 signal. The image shown is representative of the data used for quantifications shown in D. (**D**) Quantification of immunofluorescence images of ALI-CO-derived dissociated neurons labeled for five distinct ribosomal proteins. The histograms report the mean pixel grey value along axons and dendrites (mean±SD). Each data point represents a different axon or dendrite. With the exception of quantifications done on the ribosomal protein S6, axons were identified as SMI312^+^/MAP2^−^ neuronal processes, while dendrites were identified as SMI312^−^/MAP2^+^ neuronal processes. Due to antibody incompatibility, in the case of S6, dendrites were identified as MAP2^+^ processes while axons were identified as GFP^+^/MAP2^−^ processes. Mann-Whitney tests were employed for statistical analysis: RPL8, n=10, p<0.0001 (****); RPS10, n=12, p<0.0001 (****); RPS16, n=12, p<0.0001 (****); RPS26, n=12, p<0.0001 (****); S6, n=12, p=0.0001 (***). Scale bars: 50 nm in (A), 20 μm and 2 μm in (C).

## Discussion

To our knowledge, our dataset represents the first cryo-EM resource of developing human axon shafts. To date, cryo-ET studies of neurons have been conducted on rodent primary cells grown on EM grids (7, 18, 30). Such dissociated neurons exhibit a random pattern of neurites and do not form the directed long-range axon tracts observed *in vivo* in the tissue context. Additionally, the growth state of axons is not always clear. Moreover, cryo-EM of human neurons has so far remained out of reach. These limitations are overcome by the combination of human cerebral organoids with cryo-CLEM, allowing visualization of the intracellular landscape of the shafts of actively growing human axons, unambiguously identified and in an environment that closely mimics the physiological context of the developing nervous system.

Our findings reveal structural characteristics that point to architectural and molecular adaptations of growing axon shafts and can help explain their unique behaviors. The organization of the cytoskeleton, in particular the dense microtubule bundles, likely reflects the importance of mechanical properties, such as rigidity, and of cytoskeletal transport for vectorial projection of axons over long distances. Reminiscent of textbook electron micrographs of mature nervous tissue, growing axon shafts also appear to contain exclusively smooth ER. Its intricate morphology and its interaction with other membranes suggest that the axonal ER is primarily involved in lipid biosynthesis to provide membrane material. Consistent with this, we observed membrane contact sites between the ER and the plasma membrane, likely representing sites for transfer of newly synthesized lipid molecules, as lipid flux is a major function of membrane contact sites in all cell types (31, 32). Furthermore, the tight association of the ER with microtubules could reflect attachment needed to pull the ER along the extending axon. Alternatively, this arrangement could represent piercing of ER cisternae by microtubules that grow quickly along the extending axon. These structural features of axonal ER may explain how axons acquire their unique surface area-to-volume ratio during elongation through lipid supply.

Our quantification of ribosome density provides an estimate of the potential for local protein biosynthesis. Previous EM studies have reported clusters of ribosomes in the axon initial segment and growth cones (3, 33–37). Mature presynaptic termini were shown to harbor ribosomal components, consistent with the presence of a distal ribosome pool, and metabolic labeling experiments demonstrated presynaptic protein synthesis (38, 39). Within the extending axon shaft, the presence of an active pool of ribosomes is less clear. In contrast to the known hallmarks for identifying pre- and post-synaptic compartments, axonal shafts are not easily identified through defining structural features; they can be mistaken for thin dendrites and have therefore remained largely unexplored. Our experimental setup allows us to confidently ascribe our observations to axon shafts within a neuronal tissue context, and reveal that this specific region exhibits low levels of ribosomes. This observation is complementary to previous findings of local translation at growth cones and presynaptic termini, and suggests that different regions of the axon can exhibit very different molecular landscapes. We therefore propose a refined model in which biological processes in initial segments of the axon, the growth cone, and presynaptic termini rely on local translation(3, 33–39), while the axon shaft during growth is depleted of translational machinery and consequently must have limited capacity for new protein synthesis.

Our findings may have implications for how the axonal compartment becomes established and how axons navigate their environment. Our data suggest that already in the early stages of neurodevelopment, during axon pathfinding, the axon shaft differs from dendrites in that it displays reduced ribosome levels. Perhaps this early differential distribution of ribosomes is one of the factors that set the axonal and dendritic compartments apart. Furthermore, because protein synthesis in the growth cone was shown to be used primarily for navigation and not for extension (40), we speculate that ribosomal depletion along the shaft could serve as a potential mechanism to desensitize this structure to exogenous guidance cues and ensure correct wiring. The axon shaft represents the largest portion of the axon, acting as a long-distance highway connecting the soma to the growth cone first and to presynaptic compartments later, yet it is arguably the least studied part of the axon. Since it is this specific region that is damaged in nerve injury, a deeper understanding of the cell biology of the axon shaft will be highly informative.

## Materials and Methods

### Cell and cerebral organoid culture

The study employed H9 human embryonic stem cells (Wisconsin International Stem Cell Bank, Wicell Research Institute, WA09 cells) approved for use in this project by the U.K. Stem Cell Bank Steering Committee, cultured under feeder free conditions in StemFlex (Thermo Fisher Scientific, A3349401) on Matrigel (Corning, 356230) coated plates and passaged twice a week using 0.7 mM EDTA in sterile D-PBS without Ca^2+^ and Mg^2+^. Cerebral organoids were generated and grown using the STEMdiff Cerebral Organoid Kit (Stem Cell Technologies, 08570) according to manufacturer’s guidelines. The HeLa cell line was stably expressing doxycycline-inducible for Fsp27-EGFP and was grown as an adherent culture in a high-glucose DMEM media containing GlutaMAX (Thermo Fisher Scientific, 31996). The media was further supplemented with 10% Tet-approved heat-inactivated FBS (Pan Biotech p30-3602), 10 mM HEPES pH 7.2, 0.2 mg/ml Hygromycin B (Invitrogen, 10687010) and 1x non-essential amino acids solution (Thermo Fisher Scientific, 11140050).

### Plasmid construct generation

The study made use of the Sleeping Beauty transposon system. The constructs pCAGEN-SB100X, pT2-CAG-fGFP plasmid (Addgene #108714) and pT2-CAG-fFusionRed were previously used and described in other studies (*12*, *15*). EGFP-E-Syt1 was a gift from Pietro De Camilli (Addgene plasmid # 66830) (*41*). The construct pT2-CAG-GFP-E-Syt1 was generated by restriction digestion of the EGFP-E-Syt1 (Addgene #66830) and the pT2-CAG-fGFP plasmids with AgeI and MluI. The fragment encoding GFP-E-Syt1-SV40 PolyA was then ligated into the pT2-CAG backbone using T4 ligase. pcDNA3-hL1 was a gift from Fritz Rathjen (Addgene plasmid # 89411). The construct pT2-CAG-hL1CAM-GFP was generated by Gibson assembly on the fragments here described; the pT2-CAG-fGFP vector was linearised by restriction digestion with EcoRI and MluI, the fragment encoding the hL1CAM ORF was PCR amplified from the pcDNA3-hL1 plasmid (Addgene #89411) with primers Fwd: 5’-AACGTGCTGGTTATTGTGCTGTCTCATCATTTTGGCAAAGAAAGATGGTCGTGGCGCT-3’ and Rev: 5’-TACAAGAAAGCTGGGTACCGTTCTAGGGCCACGGCAGGG-3’, the fragment encoding EGFP-SV40 PolyA was PCR amplified from the pcDNA3-hL1 plasmid (Addgene #89411) with primers Fwd: 5’-ACCCTGCCGTGGCCCTAGAACGGTACCCAGCTTTCTTGTACA-3’ and Rev: 5’-CAGCCGGGGCCACTCTCATCAGGAGGGTTCAGCTTACTCAAGAGGCTCGAGTGCAACTATG-3’. The Q5 High-Fidelity 2x Master Mix (New England Biolabs, M0492S) was used for PCR. The ligation and Gibson assembly products were cloned in TOP10 chemically competent *E.coli*. Plasmids recovered from bacterial colonies were screened by restriction digestion and the correct products were verified by sequencing.

### Construct expression in organoids

Organoids were electroporated as previously described with expression constructs (*11*, *12*, *15*). Briefly, a total of 5 μl of a 1 μg/μl plasmid solution (750 ng/μl transposon donor plasmid and 250 ng/μl pCAGEN-SB100X) was injected into the ventricles of 45-55 day-old organoids and electroporated using the BTX Gemini X2 HT Electroporation System (BTX, 452008) and 5 mm gap Petri dish platinum electrode kit to deliver 5 square-wave 1 ms pulses of 80 V amplitude with 1 s inter-pulse intervals. Approximately one week after electroporation the organoids were prepared for ALI culture.

### ALI-CO preparation

Cerebral organoids aged 45-60 days were prepared for ALI culture as previously described (*15*). In brief, organoids were embedded in 3% low-gelling temperature agarose (Sigma-Aldrich, A9414) in HBSS without Ca^2+^ and Mg^2+^ (Thermo Fisher Scientific, 14175095) and sectioned into 300 μm-thick slices on a Leica VT1000 S Vibrating blade microtome. All surrounding agarose was removed from the tissue and 2-3 tissue slices were positioned on each Millicell Cell Culture Insert (Merck Millipore, PICM0RG50) using No.22 scalpels (Swann-Morton, 0508). The slices were incubated for 1-2 h in SSSC medium (0.5% glucose, 10% FBS and 1x Anti-Anti in high glucose DMEM supplemented with Glutamax) and cultured long-term in SFSC medium (1x B27, 0.5% glucose, 1x Glutamax and 1x Anti-Anti in Neurobasal medium) with daily half-media changes.

### Organoid dissociation and neuronal culture

Mature organoids aged between 50-80 days were dissociated using ACCUMAX cell dissociation reagent (Sigma Aldrich, A7089) supplemented with 400 μg/ml DNAse I. For each organoid dissociated, 0.5 ml of dissociation solution were used. Organoids were resuspended in dissociation solution and subject to 4× 5 min incubation steps in an incubator at 37° C; after the first 5 min the organoids were resuspended by flicking the tube, after 10 min the organoids displayed a fluffy appearance and were pipetted up and down once, then broken into cell clumps After 15 min the cell clumps were resuspended by pipetting up and down 3-5 times and then 10 more times after an additional 20 min incubation. Dissociation was stopped by addition of an equal volume of maturation medium (Stem Cell Technologies, 08570). The cell suspension was passed through a 70 μm nylon cell strainer (Corning, 352350). A small aliquot was taken for a live cell count and the remaining cell suspension was spun down at 300x g for 5 min. The cell pellet was resuspended in SFSC medium and 50,000 cells were seeded into each well of 8 well Lab-Tek II glass chamber slides (Nunc, 154534) for immunofluorescence preparation.

### Immunofluorescence sample preparation

Dissociated neurons were fixed in 4% PFA for 10 min at room temperature and incubated in permeabilization buffer (4% donkey serum and 0.25% Triton-X in PBS without Ca^2+^ and Mg^2+^) for one hour at room temperature prior to overnight staining with primary antibodies in blocking buffer (4% donkey serum and 0.1% Triton-X in PBS without Ca^2+^ and Mg^2+^) at room temperature. Antibodies used in this study with the corresponding dilution factor were: rabbit anti-RPL8 (Abcam, ab169538, 1:200), rabbit anti-RPS10 (Abcam, ab151550, 1:200), rabbit anti-RPS16 (Abcam, ab26159, 1:200), rabbit anti-RPS26 (Thermo Fisher Scientific, PA5-65975, 1:200), Alexa Fluor 647 conjugate mouse anti-S6 ribosomal protein (Cell Signaling Technology, 5548), mouse anti-SMI312 (BioLegend, 837904, 1:500), chicken anti-MAP2 (Abcam, ab5392, 1:500). The next day the slides were washed three times in PBS, followed by a one hour incubation at room temperature with 405, 568 and 647 Alexa Fluor conjugate secondary antibodies diluted 1:500 in blocking buffer. After secondary antibody staining, the slides were washed three times in PBS and the coverslips were mounted using ProLong Diamond antifade mountant (Thermo Fisher Scientific, P36961).

### EmGFP Sendai virus transduction and SiR-tubulin labelling

Dissociated neuronal cultures used for immunofluorescence staining of ribosomal subunits were fed with 200 μl of SFSC medium supplemented with CytoTune EmGFP Sendai Fluorescence Reporter (Thermo Fisher Scientific, A16519, 8.1×10^7^ CIU/ml) diluted 1:200. After 3-4 days the cells started displaying EmGFP signal. Approximately two weeks after dissociation, cultures produced thin EmGFP^+^ axons and the cultures were fixed for analysis. SiR-tubulin (*42*) was reconstituted in sterile DMSO to a concentration of 1 mM. For staining of ALI-COs SiR-tubulin was diluted to a final concentration of 1 μM in SFSC medium and applied dropwise to the top of the slice using a controlled oral-suction pipetting apparatus and care was taken not to disturb the grids. After approximately one hour at 37 °C and 5% CO_2_ samples were ready for imaging.

### Fluorescence image acquisition and analysis

Widefield fluorescence images were acquired on a Nikon ECLIPSE Ti2 system at 10x (0.3NA) and 20x (0.75NA) magnification and on an EVOS FL inverted microscope (Thermo Fisher Scientific). Confocal images of SiR-tubulin stained organoids were acquired on a Zeiss LSM 710 upright system at 10x (0.3NA) magnification. The time course of fGFP^+^ axon growth on grids was acquired on a Zeiss LSM 780 confocal microscope using a 10x (0.3NA) objective and a pixel size of 830 nm. Samples were incubated at 37 °C and 5% CO_2_ in 35 mm Easy-Grip tissue culture dishes (Corning, 353001). The microscope objective was aligned to the center of the grid and 4×4 tiled-images were acquired approximately every 12 hours. For measurement of axon growth rates, live FM movies of ALI-COs were acquired on a Zeiss LSM 710 and Zeiss LSM 780 inverted microscope using a 10x (0.3 NA) objective and a pixel size of 1.384 μm. Samples were incubated at 37 °C and 5% CO_2_ in 35 mm Easy-Grip tissue culture dishes (Corning, 353001) and images were acquired every 12 min. The manual tracking plugin in ImageJ was used to track the position of individual growth cones throughout the movie frames. The time interval was set to 12 min and the x/y calibration to 1.3837 μm. For immunofluorescence analysis of ribosomal subunit distribution in axons and dendrites, dissociated human neurons were imaged on a Zeiss LSM 780 confocal microscope at 60x (1.4NA oil) magnification. The main criterion for fluorescence image acquisition was the presence of both MAP2^+^/SMI312^−^ dendrites and MAP2^−^/SMI312^+^ axons within the field of view. ImageJ was used for analysis and the mean grey value of the ribosomal protein of interest along the length of two axons and two dendrites per image was calculated. Axons were identified as SMI312^+^/MAP2^−^ or, due to antibody incompatibility, in the case of S6 as GFP^+^/MAP2^−^ processes. Dendrites were identified as MAP2^+^ processes. The ribosomal proteins imaged include: RPL8, RPS10, RPS16, RPS26 and S6. For each of these targets the average mean grey value was calculated across axonal and dendritic segments and a Mann-Whitney unpaired two-tailed test was used for statistical comparison between the two groups.

### Electron cryo-microscopy (cryo-EM) sample preparation

After 4-7 days the ALI-COs started to display escaping processes and were thus considered ready for grid placement. Quantifoil R2/2 or R3.5/1 200 mesh Au grids with carbon film (Quantifoil) were coated with 0.01% poly-L-ornithine solution (Sigma Aldrich, P4957) overnight at 4°C. The next day the grids were further coated with a solution of 10-20 μg/ml Laminin (Sigma Aldrich, L2020) and 0.001% Fibronectin (Sigma Aldrich, F0895) in ultrapure water at room temperature for four hours. Organoid sections were inspected on an EVOS FL inverted microscope (Thermo Fisher Scientific) by brightfield or GFP fluorescence. In the case of unlabeled organoids the grids were placed at sites where single escaping processes could be seen by brightfield, in the case of GFP^+^ organoids the grids were placed near the fluorescent foci. Cryo-EM grids were blotted with Whatman filter paper grade 1 (GE Healthcare) and placed in direct contact with the edge of the organoid section. Importantly, the edge of the grid was juxtaposed to that of the organoid section, yet not covered by it. In the case of GFP^+^ organoids growth of the processes could be monitored daily, in the case of unlabeled organoids, after approximately two weeks SiR-tubulin (Spirochrome, CY-SC002) was applied dropwise on top of the grids to visualize axon tracts. The grids were deemed ready for freezing earliest after 10-14 days, or once the axons reached an area within 5 grid squares of the grid center. Immediately prior to freezing, the grids were hydrated by applying media dropwise using a controlled oral-suction pipetting apparatus and glass capillaries. This step was crucial to reduce desiccation of neuronal processes during blotting of the EM grids. Tracts on EM grids were separated from their cell bodies using a 3.5 mm disposable biopsy punch (Integra, 33-33) and immediately collected with an L5 clamp style thin-tip tweezer (Dumont, 72882-D), backside blotted for 5-10 s with Whatman filter paper grade 1 (GE Healthcare) and vitrified in liquid ethane using a home-made manual plunger.

### Cryo-fluorescence microscopy (cryo-FM)

The grids were screened for ice thickness and fluorescent signals of fGFP^+^ within axon tracts by cryo-FM with the Leica EM cryo-CLEM system. The system was equipped with a HCX PL APO 50x (0.9 NA) cryo-objective (Leica Microsystems), an Orca Flash 4.0 V2 SCMOS camera (Hamamatsu Photonics), a Sola Light Engine (Lumencor), a L5 filter (Leica) for detection of GFP and a Y5 filter (Leica) for the detection of the SiR-tubulin stain. The microscope stage was cooled to −195 °C and the room was humidity controlled (below 25 %). A 2.0 × 2.0 mm montage of each grid was taken of the green (1 s, 30% intensity), brightfield channel (30 ms, 70), and optionally of the far-red channel (1 s, 30% intensity). Individual z-stacks of grid squares of interest were acquired in 1 μm steps to cover the full range of fluorescent signals. Correlation of fluorescent axon tracts on cryo-EM grid square maps was done in Icy using the ec-CLEM plugin (*43*) using landmark features and carbon film holes in cryo-FM and cryo-EM images.

### Preparation and focused ion beam milling (FIB-milling) of control HeLa cells

Control HeLa cell samples were prepared as described in (*29*). In short, HeLa cells were grown for 24 h on holey carbon-covered gold EM grids (200 mesh, R2/2, Quantifoil), fed with oleic acid and induced with doxycycline for Fsp27-EGFP expression for an unrelated project. 16h post-induction, HeLa cells were stained for 1 h with LipidTOX Deep Red dye (Thermo Fisher Scientific, H34477). Subsequently, grids were manually backside blotted with Whatman filter paper No 1 and immediately vitrified in liquid ethane using a manual plunger. Thin lamellae were generated by cryo-FIB milling performed with a Scios DualBeam FIB/SEM (FEI) equipped with a Quorum stage (Quorum, PP3010T) in a procedure similar to the one described in (*44*). Prior to milling, grids were coated with organometallic platinum using a gas injection system for 30s at 13mm working distance and 25° stage tilt. The electron beam was used for locating the cells of interest at 5 kV and 30 pA beam current and for imaging to check progression of milling at 2 kV and 30 pA beam current. Milling was performed with stepwise reduction of the ion beam current (from 30 kV, 1 nA to 16 kV, 23 pA) while changing the stage tilt as described in (*45*). The lamellae with a 10° pre-tilt were milled to a final thickness below 300 nm.

### Electron cryo-microscopy (cryo-EM)

Cryo-EM grids were screened on a Tecnai T12 (FEI) with an Orius camera or a Tecnai F20 (FEI) with a Falcon 2 detector (FEI) by mapping the central parts of the grids using SerialEM (*46*) at pixel sizes of 132 nm or 87 nm, respectively. The preservation of axon tracts on individual grids squares was examined on images acquired at pixel sizes of 6.3 nm or 6.0 nm, respectively. Cryo-ET data acquisition was done using SerialEM on a Titan Krios microscope (Thermo Fisher) equipped with a Quantum energy filter and a K2 direct electron detector (Gatan) operated in counting mode. Montaged images of the central part of the grid were acquired in linear mode with 171 nm pixel size. Montages of individual grid squares with axon tracts or lamellae of HeLa cells were taken with 5.1 nm pixel size. These montages were used for correlation to fluorescent axon tracts, based on landmark features with the Icy software using the ec-CLEM plugin (*43*). Tilt series were acquired at areas of interest in low-dose mode from 0° to ±60° using a grouped dose-symmetric tilt scheme with 1° increment, a group size of 4 (*47*), and a pixel size of 3.7 Å; both for the axon tracts and the control HeLa cells. The target dose rate was kept around 4 e^−^/px/s on the detector. The energy filter slit width was set to 20 eV. Tilt images were acquired as 4 frames with approximately 1 e^−^/A^2^ dose per tilt image. The nominal defocus for all tilt series was set to −5 μm. The frames of tilt series images were aligned with IMOD alignframes. The tilt series were aligned in IMOD using patch tracking and then reconstructed at a pixel size of 7.4 Å as backprojection tomograms with SIRT-like filter corresponding to 10 iterations (*48*, *49*). To improve visibility for representation in figures, gaussian filtering was applied to the shown tomographic slices. For analyzing the microtubule polarity by subtomogram averaging, the contrast transfer function was estimated and corrected for by phase flipping in IMOD, and the tomograms were reconstructed by unfiltered backprojection at 7.4 Å pixel size. Five of the 9 HeLa tomograms used here have been analyzed and published before (*29*).

### Image processing and analysis

The segmentation model shown in Fig. 2B and in Supplemental Movie 2 was generated using Amira (Thermo Fisher Scientific) and IMOD (Kremer et al. 1996). Membrane surfaces were segmented manually, followed by extensive smoothening and simplification. Microtubules and actin filaments were first modelled as tubes in IMOD and then imported as tubular volumes into Amira. Within the segmented volumes of individual membrane objects and cytoskeleton objects, grey value-thresholding was used for a second segmentation step to eventually depict only high-density voxels within the objects of interest. Note that the segmentation model is inverted along the z-axis relative to the original tomogram. Movie S3 was generated by first selecting a region of interest from tomogram EM, followed by identifying and aligning images obtained from the previous successive imaging steps of fluorescent live imaging, cryo-FM, and electron tomography. Each acquired image was then used to generate a stack of progressive magnification into the region of interest using an ImageJ macro (zoom_movie_ImageJ_v2) written by Eugene Katrukha and provided through GitHub Gist. All stacks were then converted and concatenated into a single movie.

The microtubule directionality was determined from the radial tilt of protofilaments in averages from individual microtubules. Points along individual microtubules were clicked along their center to create a contour. Overlapping subvolumes were extracted along each contour and averaged together using a cylindrical mask and one individual subtomogram as a reference in PEET (*50*, *51*). Microtubule directionality was determined from axial views of individual subtomogram averages (*24*). To measure the plasma membrane surface area of axon shafts, the plasma membrane boundaries of individual axons were segmented in tomograms at bin 10 that were rotated around the x-axis. Axons were only analyzed if they were fully contained in the tomogram volume. Segmentations and surface area measurements were performed in IMOD. Each measurement was normalized to the individual axon length. The ER diameter of the thinnest ER tubules contained in the axon and HeLa datasets were measured in IMOD as the distance between two points on the cytosolic edge of the membrane of each ER tubule. Significance was tested with Welch’s t test in Graphpad Prism.

Ribosome-like particles were identified visually on the basis of their size, shape and contrast in the axon tomograms, in reference to other tomograms of eukaryotic cells that contained polysomes and monosomes. To better visualize the topographic distribution of the ribosome-like particles, example volumes of 0.05 μm^3^ from cytosolic areas of the tomograms are shown in Figure 4A, in which the position of each ribosome-like particle is illustrated by a sphere with 30 nm diameter. ‘Other processes’ comprised 4 tomograms containing high numbers of ribosomes. These tomograms were acquired at the edge of the EM grid, which is in close proximity to the cell bodies of the organoid slice. Additionally, two of these tomograms were acquired at the ending of a process. For these reasons, we interpreted them as representing cellular processes other than axonal tracts, excluded these 4 tomograms from the data on axon shafts and classified them as ‘other processes’. The number of ribosome-like particles found within each tomogram was normalized to the tomographic volume. The tomographic volume was estimated from the x, y dimensions of tomograms. The z dimensions were estimated at the tomogram center. Using Avogadro’s number, the average molar concentration of ribosome-like particles within the tomograms of the axon tracts was calculated. Due to the permissive assignment and quantification accounting for their scarcity, it cannot be excluded that some of the ribosome-like particles identified in the axon shaft may actually correspond to other protein complexes. Thus, the calculated concentration in axon shafts is likely an overestimate.

## Author contributions

S.L.G. and P.C.H. conceived the project, designed and conducted experiments, analyzed data, and wrote the manuscript. I.G. conducted experiments on HeLa cells. M.R.W. and M.S. contributed to and assisted with experiments. M.A.L. and W.K. conceived and supervised the project, analyzed data, and wrote the manuscript.

## Acknowledgments

The authors would like to thank members of the Kukulski and Lancaster labs for helpful comments and discussion, the light microscopy and EM facilities of the MRC Laboratory of Molecular Biology for support during data acquisition, Paul Donlin-Asp, Erin M. Schuman and Benoît Zuber for critical reading of the manuscript. Work in the Lancaster lab is supported by the Medical Research Council (MC_UP_1201/9) and the European Research Council (ERC STG 757710). Work in the Kukulski lab was supported by the Medical Research Council (MC_UP_1201/8). M.R.W. is supported in part by the Natural Sciences and Engineering Research Council (NSERC) of Canada (PGSD).

## Supplementary Figures

**Fig. S1.**
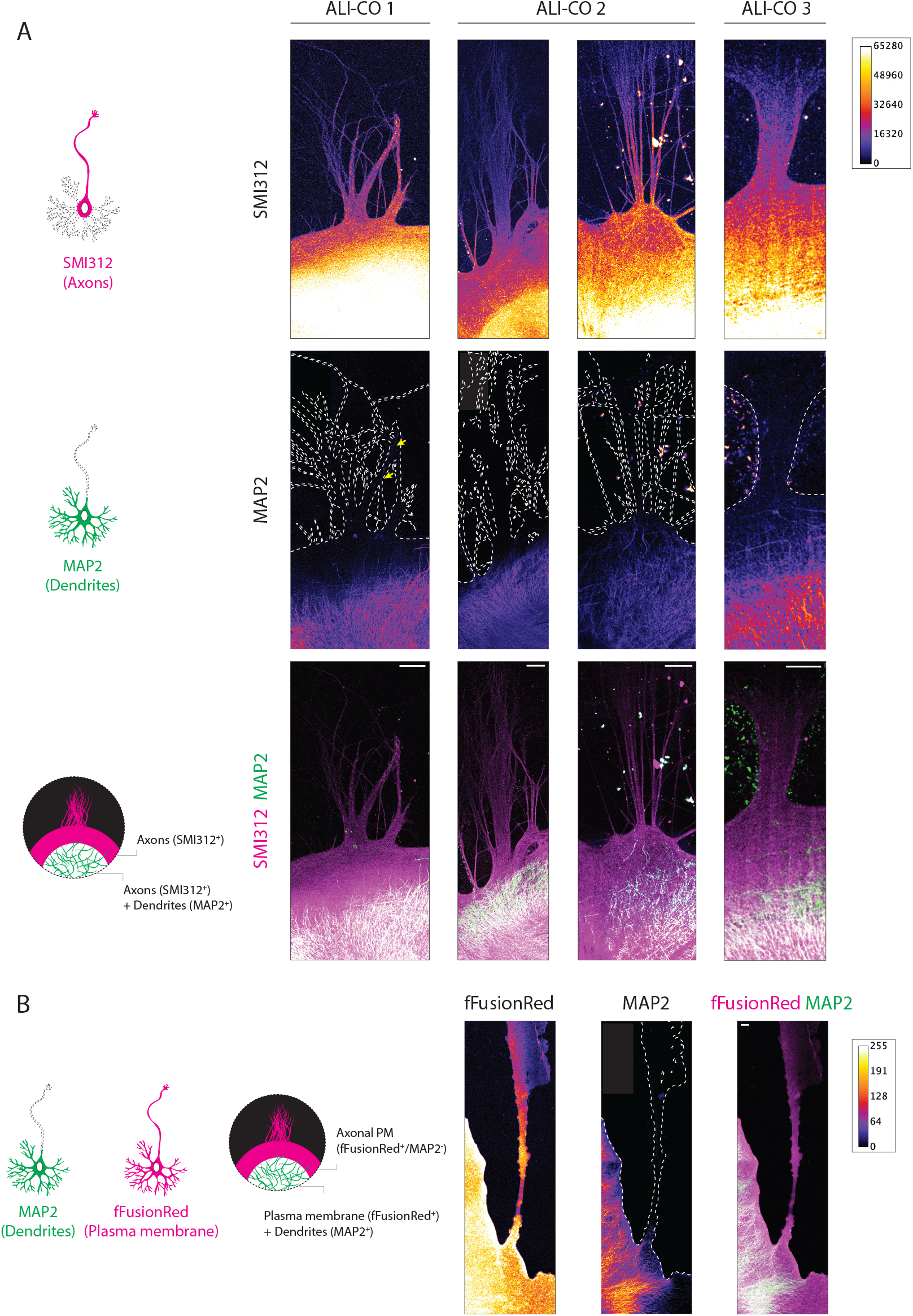
ALI-CO tracts comprise exclusively axons and lack dendrites. (**A**) Schematic representation of the distribution of SMI312 and MAP2 IF signal across a neuron and expected staining pattern in the case where dendrites are absent from exiting bundles (left). Representative images of 4 bundles from 3 different organoids stained for the panaxonal marker SMI312 and the dendritic marker MAP2, single channel heatmaps are reported in the top two rows and the bottom row reports the composite image (right). The outline of the SMI312 signal is reported on the MAP2 channel marking the surface area of the bundle. The yellow arrows in the left MAP2 panel point to a single spurious neuronal cell body within the axon bundle. The heatmap reports gray value signal intensity ranging 0-65280. (**B**) Schematic representation of the distribution of fFusionRed and MAP2 IF signal across a neuron and expected staining pattern in the case where dendrites are absent from exiting bundles (left). Representative image of an escaping bundle from one ALI-CO stained for the plasma membrane marker fFusionRed and the dendritic marker MAP2. The outline of the fFusionRed signal is reported on the MAP2 channel marking the surface area of the tract. The heatmap reports the gray value signal intensity. Scale bars: 100 μm in (A) and (B).

**Fig. S2.**
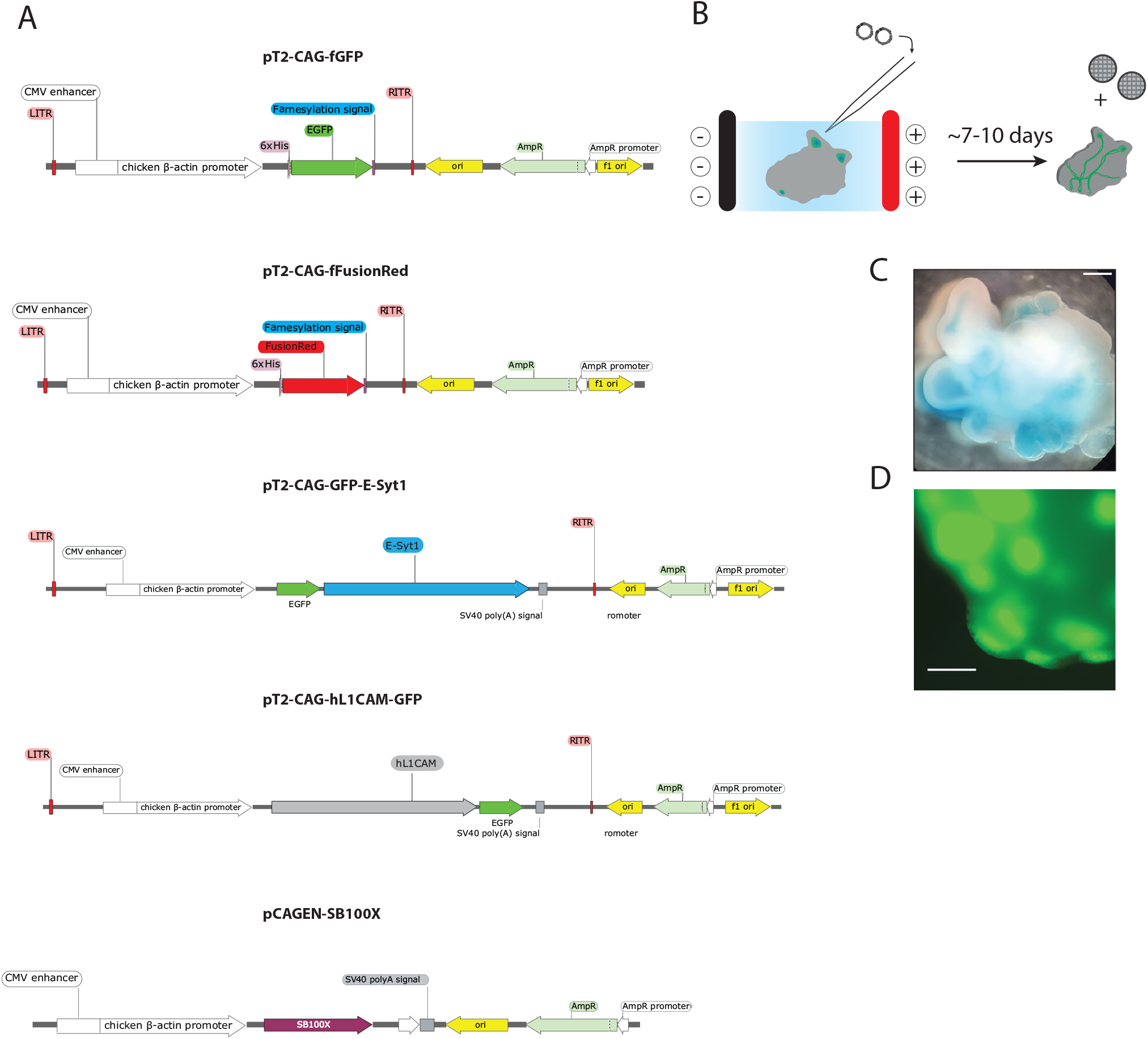
Plasmid constructs and electroporation strategy employed in the study. (**A**) Reported are the linear schematics of the plasmid maps of the constructs employed in the study. The SleepingBeauty system was used for transgene expression of the following open reading frames (ORFs) driven by CAG: farnesyl-GFP (fGFP), farnesyl-FusionRed (fFusionRed), C-terminally GFP-tagged human L1 cell adhesion molecule (L1CAM-GFP) and N-terminally GFP-tagged ESYT1 (GFP-ESYT1). Reported also is the plasmid map for the construct encoding the SB100X transposase under the control of the CAG promoter. (**B**) Electroporation schematic depicting injection of the plasmid solution into ventricles of organoids and electroporation into the progenitor cells in the ventricular zone. After approximately 4-5 days organoids were prepared for ALI culture and following an additional 4-5 day period at the ALI, EM grids were placed in close proximity to the organoid slices. (**C**) Representative brightfield image of a cerebral organoid after injection of the Fast Green-containing plasmid solution into the ventricles. (**D**) GFP signal after micro-injection and electroporation of the membrane-targeted farnesylated GFP (fGFP) construct prior to sectioning. Scale bar: 1 mm (C) and 0.5 mm (D).

**Fig. S3.**
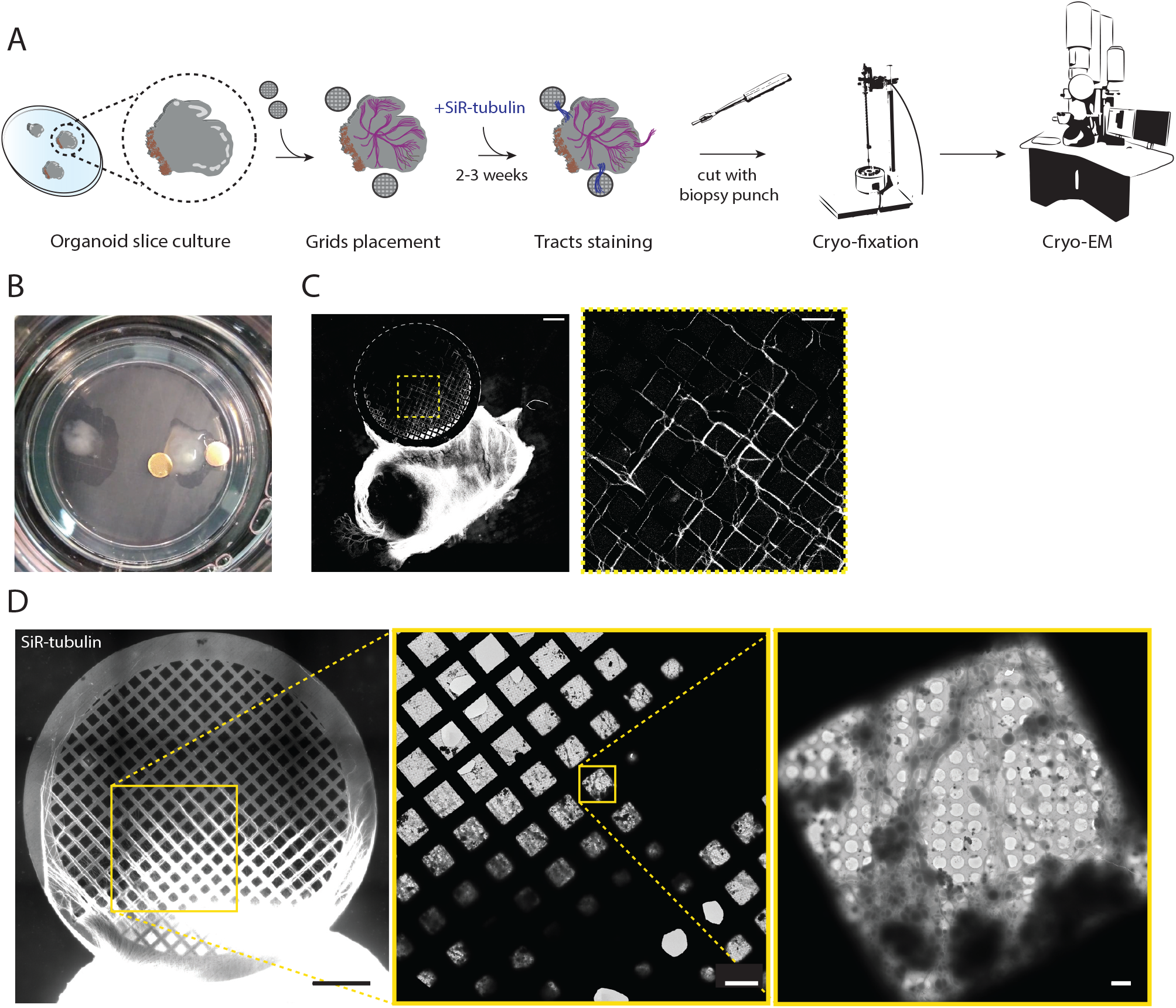
Preparation of axonal tracts from human cerebral organoid slices for cryo-EM. (**A**) Workflow for growing axonal tracts on EM grids, monitoring growth and preparation for cryo-EM. (**B**) Overview of a filter after the grids were placed adjacent to an ALI-CO. (**C**) Fluorescence microscopy overview of an ALI-CO and tracts projecting on a grid stained with the SiR-tubulin live-cell dye to monitor growth, the yellow dashed box corresponds to the area shown in magnified view on the right. (**D**) Fluorescence microscopy of tracts stained with SiR-tubulin prior to cryo-fixation (left panel). Cryo-EM overview of the outlined area on the same grid (middle panel). Cryo-EM overview of one individual grid square with axon tracts (right panel). Scale bars: 500 μm and 100 μm in (C) and 500 μm, 100 μm and 5 μm in (D) (left to right).

**Fig. S4.**
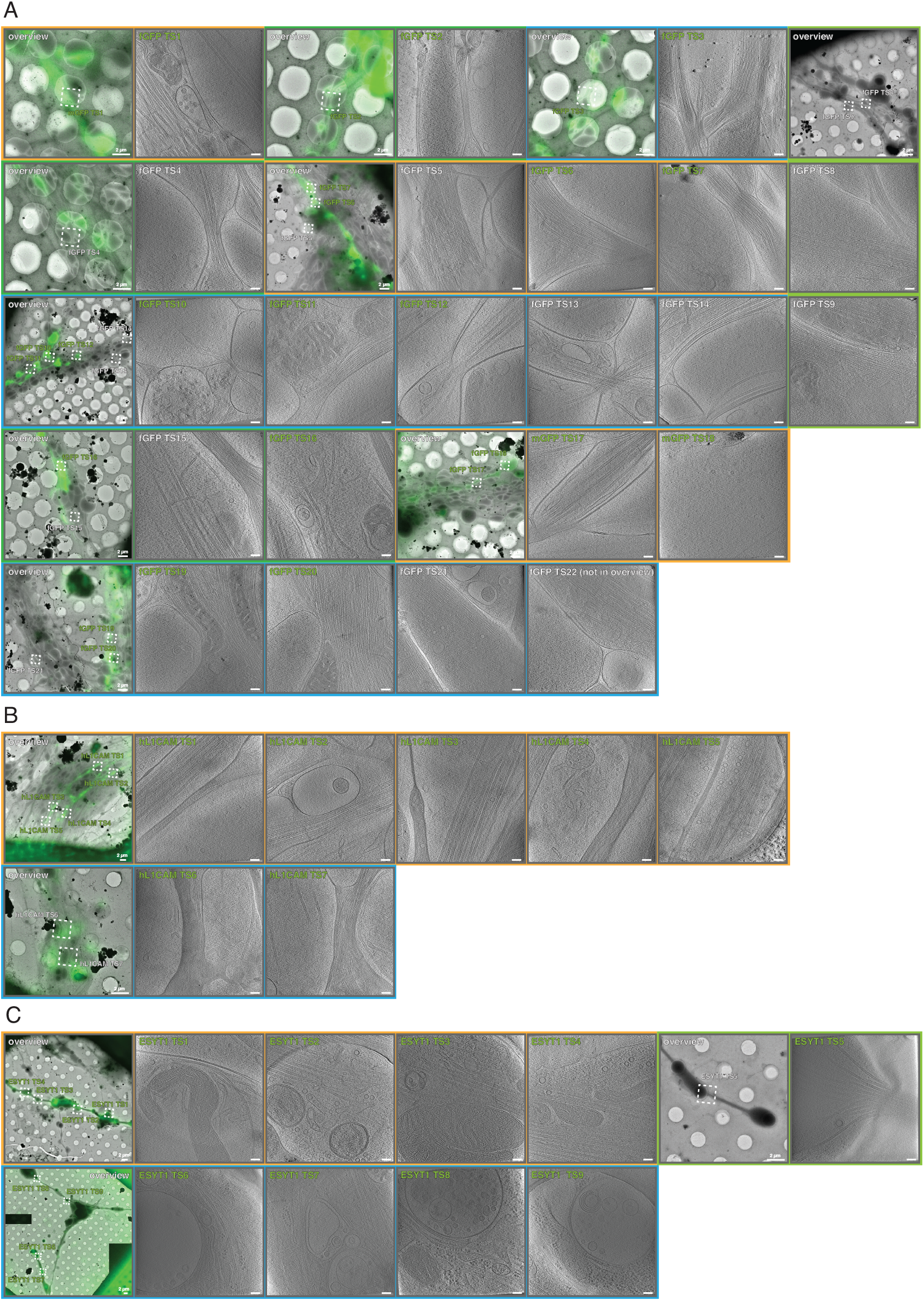
Gallery of correlative cryo-FM and cryo-ET dataset from membrane targeted GFP (fGFP), L1CAM-GFP and GFP-ESYT1 expressing ALI-COs. (**A**) Dataset of 22 axon cryo-tomograms from fGFP expressing organoids from 3 different grid preparations. Overlays between cryo-FM and cryo-EM overviews show all positions of tilt series acquisition within the axon tracts. For all positions, a corresponding virtual slice through the cryo-tomogram is shown. (**B**) Dataset of 7 axon cryo-tomograms from L1CAM-GFP expressing organoids from 2 different grid preparations. Overlays between cryo-FM and cryo-EM and corresponding virtual slices through cryo-tomograms are shown. (**C**) Dataset of 9 cryo-tomograms from GFP-ESYT1 expressing organoids from 2 different grid preparations. Overlays between cryo-FM and cryo-EM and corresponding virtual slices through cryo-tomograms are shown. The cryo-tomograms ESYT1 TS 6–9 contained ribosome-like particles. They were positioned at the outer edge of the grid, in closer proximity to cell bodies than other tracts, but also close to a growing end (lower left overview). Therefore, these tomograms likely do not represent axon shafts, and were counted as a separate class of other cellular processes, likely dendrites or axon initial segments. Scale bars: 2 μm in overlays, 100 nm in virtual tomographic slices.

**Fig. S5.**
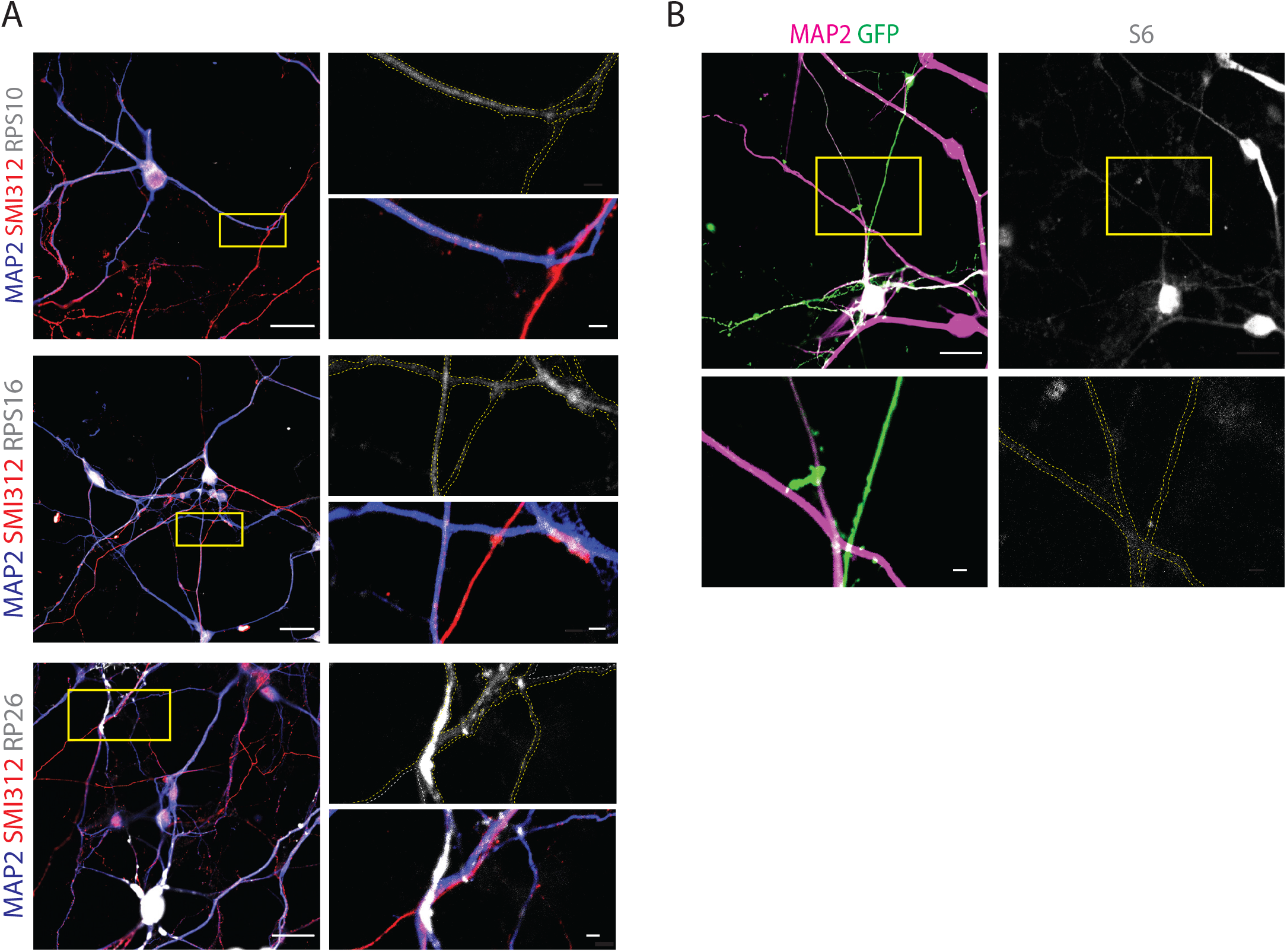
Immunofluorescence analysis of axons and dendrites supports ribosomal depletion observed by cryo-ET in the axon shaft. (**A**) Representative immunofluorescence images of dissociated organoid-derived neurons stained for the dendritic marker MAP2, the pan-axonal marker SMI312 and the ribosomal components RPS10, RPS16 and RPS26 used for the quantifications reported in Fig. 4D. Boxed in yellow is the region shown at higher magnification on the right panel as the individual ribosomal stain and the composite image. Dashed yellow is the outline of the processes reported on the single channel image of the ribosomal stain. The data highlights how dendrites (MAP2^+^/SMI312^−^) are enriched in ribosomal components compared to axons (MAP2^−^/SMI312^+^) of comparable thickness. (**B**) Representative immunofluorescence images of dissociated organoid-derived neurons transduced with Sendai EmGFP and stained for the dendritic marker MAP2 and for the ribosomal components S6. Due to antibody incompatibility, in this case dendrites were identified as MAP2^+^ processes while axons were identified as GFP^+^/MAP2^−^ processes. These images were used for the quantifications reported in Fig. 4D. Boxed in yellow is the region shown at higher magnification in the bottom panel as the S6 stain (right) and the MAP2-GFP composite stain (left). Dashed in yellow is the outline of the processes reported on the single channel image of the S6 stain. The data highlights how dendrites (MAP2^+^) are enriched in ribosomal components compared to axons (GFP^+^/MAP2^−^) of similar thickness. Scale bars: 20 μm in the overview images in (A) and (B), 2 μm in the high-magnification inserts.

---

**Movie S1:** Time-lapse movie of the extending fGFP^+^ axon shown in Fig. 1B. Images were acquired at 12 min intervals for a total duration of 13 hr. The progressive blue line marks the trajectory of the axon over the course of the movie. The axon shown corresponds to axon number 4 in the quantifications reported in Fig 1C. Scale bar: 50 μm.

**Movie S2:** Time-lapse movie over 10 days of fGFP^+^ axon tracts growing on an EM grid. Growth was monitored twice a day with approximately 12 h intervals until tracts reached the center of the EM grid. The same grid is shown in Fig 1E. Scale bar: 500 μm.

**Movie S3:** Correlative microscopy of axons from growing tracts across scales to the supramolecular architecture of individual axon shafts. The movie first shows the live FM of fGFP^+^ growing axon tracts corresponding to Movie S2, then a merge of the fGFP^+^ axons (green) and all other axon tracts labeled by SiR-tubulin (magenta), taken shortly before cryo-fixation. After cryo-fixation the grid was imaged by cryo-FM, which was correlated and overlayed onto cryo-EM overviews of individual grid squares to identify individual fGFP^+^ axons. Finally, cryo-ET along axon shafts reveals their supramolecular architecture, including vesicles containing membrane protein cargo. The FM of the grid, cryo-FM and EM images of the individual grid square are shown in Fig. 1E-G.

**Movie S4:** Tomogram and segmentation of the fGFP^+^ axon shaft shown in Fig. 2B. The segmentation reveals the complex array of microtubules (magenta) and their intimate association with the ER (cyan), which forms thin tubules that are wrapped around individual microtubules. Other highlighted cellular components include actin filaments (orange), mitochondria (green), small vesicles (yellow) and the plasma membrane of the axon shaft (light blue) and of neighboring axons (dark blue).

